# Sex-specific population dynamics and demography of capercaillie (*Tetrao urogallus* L.) in a patchy environment

**DOI:** 10.1101/576876

**Authors:** Ben C. Augustine, Marc Kéry, Juanita Olano Marin, Pierre Mollet, Gilberto Pasinelli, Chris Sutherland

## Abstract

1. Modeling the population dynamics of patchily distributed species is a challenge, particularly when inference must be based on incomplete and small data sets such as those from most species of conservation concern. Open population spatial capture-recapture (SCR) models are ideally suited to quantify population trends, but have seen only limited use since their introduction.
2. To investigate population trend and sex-specific population dynamics, we applied an open SCR model to a capercaillie (*Tetrao urogallus*) population in Switzerland living in eight distinct forest patches totalling 22 km^2^ within a region of 908 km^2^. The population was surveyed using genetic sampling of scat in 2009, 2012 and 2015. We fit an open SCR model with sex-specific detection and population dynamics parameters while accounting for the patchy distribution of habitat and the uncertainty introduced by observing the population in three years only.
3. Between 2009 and 2015, a total of 143 males, 112 females and 4 individuals of uncertain sex were detected. The annual per capita recruitment rate was estimated at 0.115 (SE 0.0144) for males and 0.127 (0.0168) for females. The estimated annual survival probability for males was 0.758 (0.0241) and 0.707 (0.0356) for females. The population trajectory implied by these survival and recruitment estimates was a decline of 2% per year; however, the sex specificity of the model revealed a decline in the male population only, with no evidence of decline in the female population. Further, the population decline observed in males was explained by the demography of just two of the eight patches.
4. Using a customized open population SCR model, we determined that the endangered capercaillie in our Swiss study area had a stable female population and a declining male population, with the male decline due to population dynamics in a subset of the study area. Our study highlights the flexibility of open population SCR models for assessing population trajectories through time and across space and emphasizes the desirability of estimating sex-stratified population trends especially in species of conservation concern.

## Introduction

Estimating demographic parameters, such as population size and demographic rates, is central to ecology, conservation, and management (Kéry & Schaub 2012). This endeavour is challenging since many species are rare and difficult to observe, especially those most in need of conservation, e.g., many carnivores (Karanth & Nichols 1998, Gardner et al. 2010), fish (Danancher et al. 2004, Raabe et al. 2014) or elusive forest birds (Suwanrat et al. 2015, Mollet et al., 2015). Camera-traps (O’Connell et al., 2011), and more recently, non-invasive genetic sampling (Taberlet & Luikart, 1999) improve our ability to detect elusive species, but small sample sizes still make reliable estimation of demographic parameters difficult in practice. Recently developed spatial capture-recapture methods (SCR: Borchers & Efford, 2008; Royle & Young, 2008) provide a means to better utilise sparse data by utilizing the spatial locations of samples and have been widely adopted over the last decade (Royle et al. 2018). However, most SCR applications to date are closed population models that estimate population size and density, but not demographic rates, which are critical for quantifying population trends through time and identifying their demographic drivers.

Open population models consider parameters associated with population changes through time, such as survival and per capita recruitment. Spatially explicit versions of these models are relatively new (but see Chandler & Clark (2014), Gardner et al. (2010), Ergon & Gardner (2014), Schaub & Royle (2014) Key to the most recent open population SCR models is a model for animal relocation between primary periods—the unit of time between which demographic change is assumed to occur, typically years. An explicit model for animal relocation allows for the processes of immigration and emigration to be separated from the processes of survival and recruitment (Ergon & Gardner 2014); however, unbiased estimates of all parameters, particularly survival and recruitment, rely on the use of accurate models of individual relocation between primary periods (Ergon & Garnder 2014; Garnder et al. 2018). Most published open population SCR studies have dealt with, or at least assumed, a continuous habitat within which individuals can relocate between primary periods without constraint, but suitable habitat for many species has become highly patchily distributed. Habitat fragmentation poses problems for the conceptualization and implementation of open population SCR models developed to date, because movement may become highly constrained by the size, shape, and connectivity of the available habitat patches within a matrix of more or less unsuitable area.

A second challenge for open population models is that of an incomplete observation of the ecological process, e.g., when surveying a population over non-consecutive primary periods. These missing primary periods require a scaling of the population dynamics parameters by the period between sampling years, or alternatively, direct estimation of the latent population dynamics parameters. Maximum likelihood-based methods for open population models typically deal with missing primary periods by re-scaling the population dynamics parameters by the time between primary periods (Cooch & White, 2012). This method works for survival, e.g., the Cormack-Jolly-Seber model (Chapter 4 in Cooch & White); however, it cannot correct the estimates of parameters with more complicated dependence structures such as the availability for capture state transition parameters of a robust design model (Chapter 15 in Cooch & White, 2012). Alternatively, Bayesian methods, particularly hierarchical models using data augmentation (e.g., Royle & Dorazio, 2012), can produce posterior distributions for latent population dynamics parameters that incorporate the uncertainty introduced by missing primary period data by characterizing the probabilities of traversing all possible state transitions (e.g., Chandler & Clark, 2014).

In this paper, we present a Bayesian open population SCR model to estimate sex-specific population trends, survival, and recruitment of the capercaillie (*Tetrao urogallus*), an elusive and endangered old-growth forest obligate, in a fragmented population in Switzerland. The capercaillie poses several challenges to the application of open population modeling including the two described above. First, capercaillie habitat in most parts of Europe has become fragmented (Helle et al. 1994, Storch 2001), preventing direct application of existing open SCR models that assume continuous habitat. Second, genetic capture-recapture surveys were only conducted in 2009, 2012, and 2015, i.e., in non-consecutive years. Third, population dynamics may vary between patches as they are of differing sizes and habitat quality. And fourth, extreme sexual size dimorphism may require stratification of the model by sex to account for sex-specific detectability and population dynamics. We used custom MCMC algorithms from the **R** package *OpenPopSCR* (Augustine 2018) to estimate sex-specific population dynamics parameters while accommodating all of these challenges, producing the first estimates of survival, per capita recruitment, population growth rate and trajectory for the endangered capercaillie.

## Materials and Methods

### Study species

The capercaillie has become a conservation flagship in Switzerland and elsewhere in Europe due to its sustained population declines (Mollet et al. 2003, Jacob et al. 2010, Mollet et al. 2015) and its status as an umbrella species for other forest birds (Suter et al. 2002, Pakkala et al. 2003). While the current consensus is that capercaillie populations in Switzerland are declining, the evidence for this decline comes from data collected between 1968 and 2003 and form lek surveys that likely underestimated population sizes (Jacob et al. 2010). More recently, Mollet et al. (2015) used SCR to more reliably estimate density using genetic capture recapture; however, no estimates of population dynamics parameters such as survival, per capita recruitment, population growth rate, or trends are available from anywhere in its range.

### Study area

Our study area is located in East-Central Switzerland (Fig. 1), on the northern slope of the Alps, and consists of several mid-elevation, mostly wooded, foothill ranges up to 1,600 m above sea level, separated from each other by deep valleys with agricultural land, towns and some small lakes. The forests on the hills are dominated by Norway spruce (*Picea abies* L.) and are interspersed with many small mires and some pastures used for grazing cattle and sheep. Mountain pine (*Pinus mugo* Turra) is an important species in the tree layer in some forests, notably on boggy soils.

**Figure. 1.**
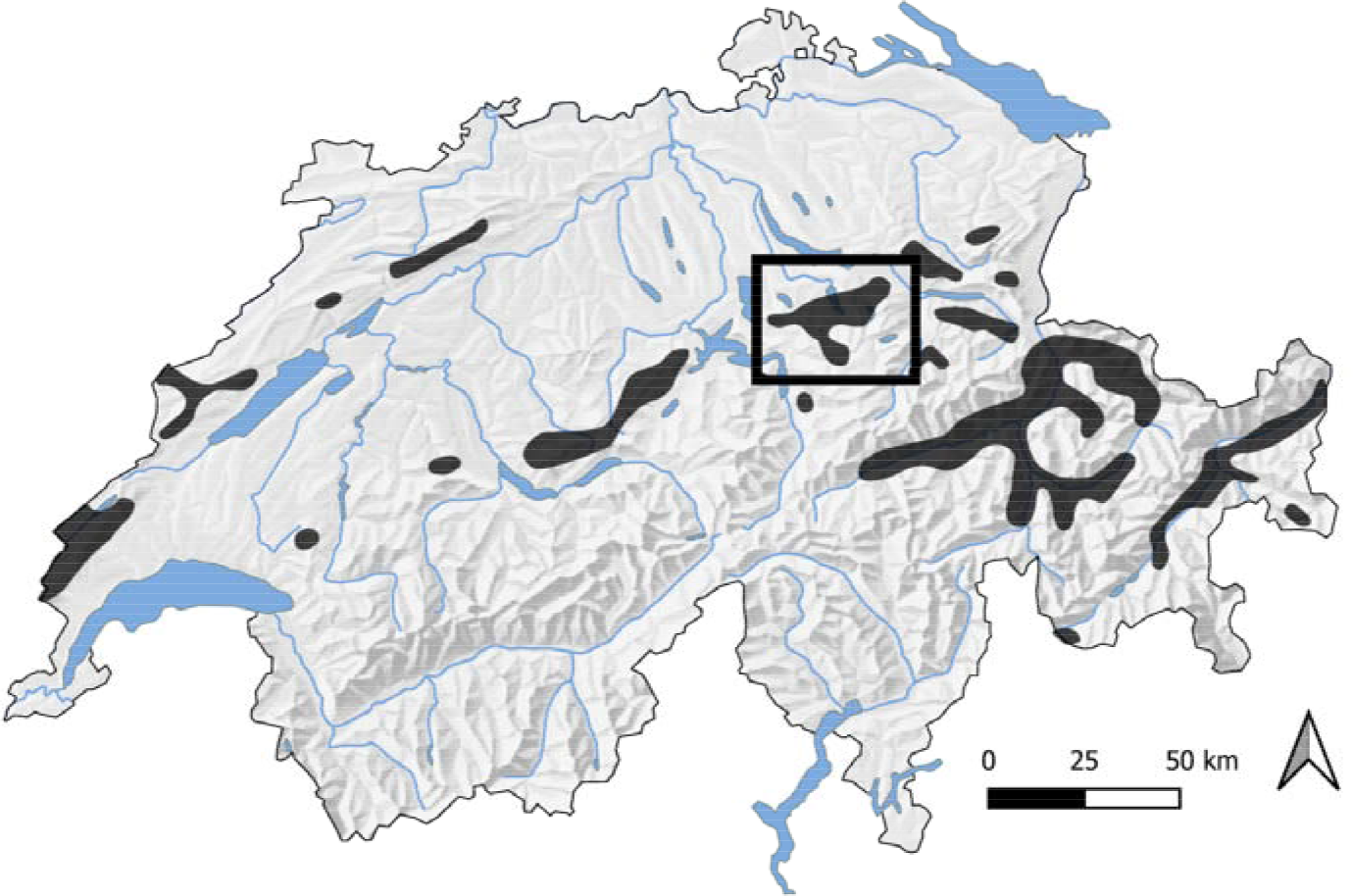
Map of the current distribution of the Capercaillie in Switzerland and location of the study area in the Canton Schwyz.

The distribution of the capercaillie in the study area is well-known due to regular surveys carried out by dedicated local bird watchers, gamekeepers, and foresters. Between 1978 and 2009, more than 1,200 opportunistic capercaillie observations were made and recorded in the database of the Swiss Ornithological Institute and in data collections of local experts. From these observations we determined that the capercaillie inhabit eight distinct patches separated by 2-6 km (Fig. 2). We defined the capercaillie habitat as follows: we generated a 500 m buffer each opportunistic observation and dissolved their borders to create a single polygon. Single observation points with buffers that did not touch any other buffer were considered isolated observations of non-resident birds and were discarded. We then intersected the observation buffer polygon with the Swiss Federal Topographic Service forest surface layer. The resulting 21.92 km^2^ area, consisting of eight fragments, was used in this study. Given that the next nearest capercaillie populations were approximately 30 km to the West, 20 km to the South and >10 km to the North-East, it was safe to assume that the focal capercaille population was geographically and demographically isolated from neighbouring populations.

**Figure. 2.**
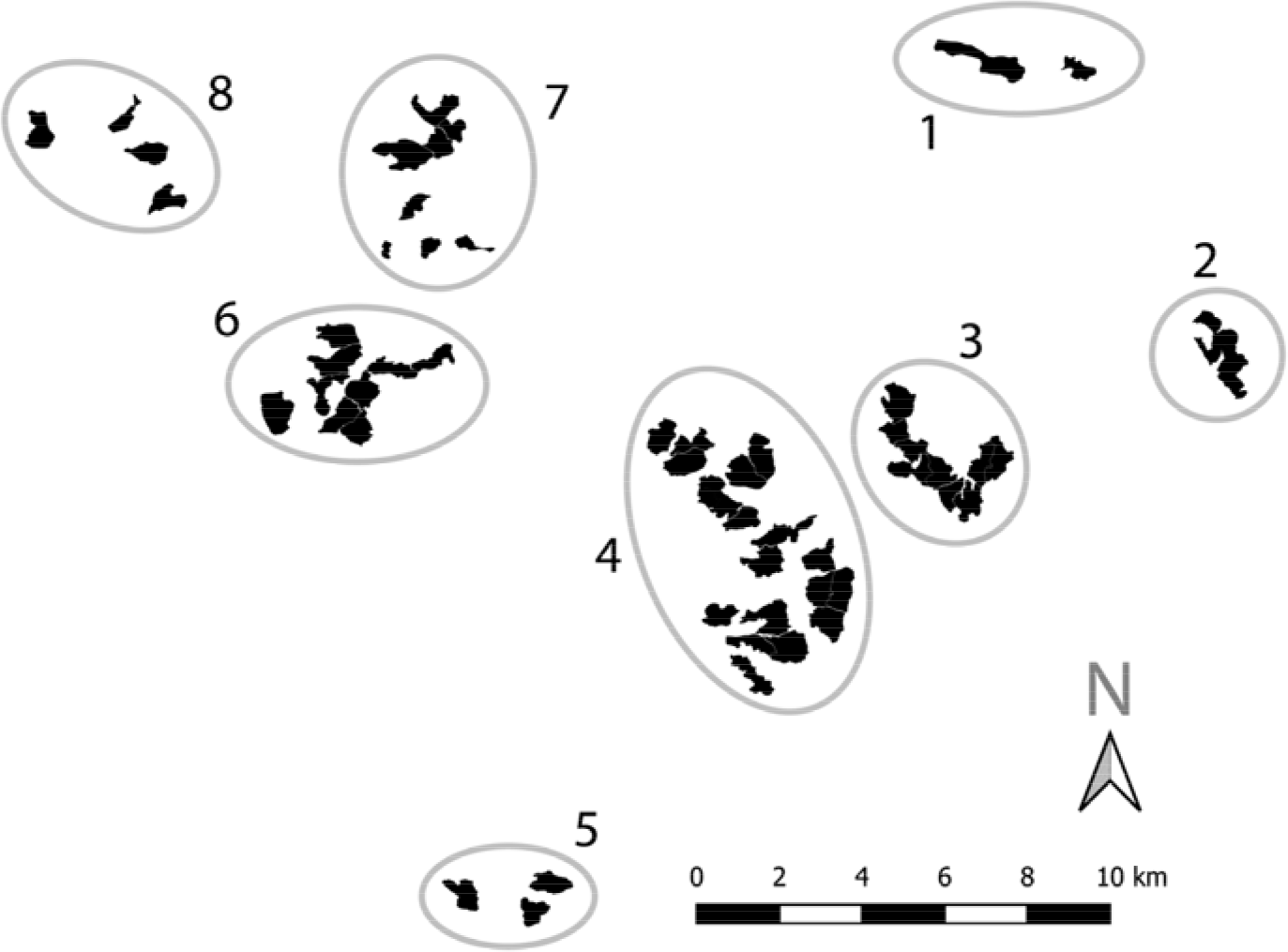
Capercaillie study area in the Swiss Canton Schwyz. A total of eight patches contain the only suitable habitat of the species within the study area, which is separated from the nearest occupied patches to the West, South, and Northeast by approximately 30, 20, and 10 kilometers.

### Data collection

Trained volunteers collected capercaillie scat samples across the study area in 2009, 2012, and 2015. Scat surveys were collected when the entire study area was completely covered with snow (mid-March to mid-April), except for Patch 6 where access was only possible from mid-April. To organize the scat collection, we sub-divided the eight patches (Fig. 2) into 58 ‘sampling, each encompassing between 20 and 80 ha (mean of 40 ha). One volunteer surveyed each unit for 6 - 8 hours. Within each unit, volunteers focused their search for scat below roosting and feeding trees, in refuge cover, along internal forest edges, around root plates, and on tree stumps, all of which are commonly used by capercaillie at a small spatial scale (Jacob et al. 2010; Bollmann et al. 2005).

For sampling units with too many suitable microhabitat features to be searched, volunteers investigated every second or third identified suitable area. Wherever scat samples or clusters of scat samples were located, the freshest single sample. Sometimes, presumably in the vicinity of lekking sites, hundreds of scat samples were clustered in small areas of 2–3 hectares. In this situation, a fresh scat sample was collected every 25 m to minimize the number of collected samples from the same individual in the same location. For each scat sample collected, the exact location was recorded with GPS (Garmin eTrex 20, accuracy approx. 5-10 m). Scat samples were stored in 50 ml plastic tubes and deep frozen at −25°C on the day of collection. We refer to Supplement A for details on DNA extraction, genotyping and population genetic analyses.

### Spatial capture-recapture models for open populations

We used a hierarchical, sex-specific, open population SCR model implemented in the **R** package *OpenPopSCR* (Augustine 2017) to estimate annual survival probabilities, per capita recruitment rates, and population growth rates of the capercaillie in our study area over the 7-year study period, while accounting for home range relocation between primary periods, and non-consecutive primary sampling periods.

*Data structure*—Our data consisted of trap-level counts of each identified individual across multiple years (primary periods). While the scat locations were recorded in continuous space, to facilitate model fitting by reducing the number of “traps” from each scat location to a smaller number of locations, we discretized the available habitat within each patch to units of approximately 14 hectares on average and associated each scat location to the nearest centroid of these discrete patch units (conceptual traps: Russell et al. 2012). This process created *J* = 153 traps, which did not vary by year, with coordinates ***X.*** The data were then formatted into an *n* × *J* × *T* 3-dimensional array, where *n* is the total number of individuals captured over the 7 years, and *T* is the number of years. Finally, following Royle et al. (2015), we introduce the potentially partially-observed 1 × *n* vector ***C*** to record individual sex with entries 0 if the observed sex is male, 1 if female, and recorded as a missing value if sex could not be determined.

*Process model*—We used the density independent population growth model described by Chandler & Clark (2014), modified for sex-specificity of the population growth rates and detection function parameters. The sex-specific population growth parameters are the male- and female-specific survival (*φ*^*m*^, *φ*^*f*^) and per capita recruitment rates (no. recruits/N; *γ*^*m*^, *γ*^*m*^). The parameters *N*_1_, *N*^*m*^_1_, and *N*^*f*^_1_ are the total, male, and female abundances in year 1, respectively. The sex-specific population abundances in years 2,…, *T* are then determined by the difference equation

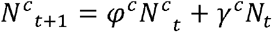

for *c* ∈ (*m*,*f*), where *N*_*t*_ and *N*^*c*^_*t*_ are the total and sex-specific abundance in year *t*, respectively. Note that sex-specific per capita recruitment is a function of the total population size, not the sex-specific population sizes. The total population size in each year is the sum of the sex-specific estimates, and population densities are the abundances divided by the area in which the individuals lived (i.e., the state space defined below).

Associated with each individual are year-specific activity centers, *s*_*il*_, which may change locations between years. To facilitate the implementation of the MCMC algorithm on this patchy landscape, the available habitat was discretised into 0.1 × 0.1 km (= 1 hectare) grid cells for a total of *G* = 2192 state space points. Then, as a discrete analogue to the typical continuous spatial uniformity assumption, we assumed 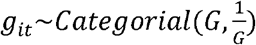 where *g*_*it*_ is the pixel identity of individuals i’s activity center in year *t*. *s*_*it*_ is then the spatial location associated with *g_it_.* Due to the extreme patchiness of the landscape and very few observed movements between the eight major patches (only 2 observed patch transitions out of 85 opportunities from individuals observed in multiple years) we assumed individuals could relocate randomly between years only within the patch in which they lived, except for the 2 individuals observed to transition patches. For the two observed individuals that did transition between patches, we assumed they transitioned in the years they were first documented in a new patch and if not observed after that, we assumed they either remained there for the remainder of the study and evaded detection or died (exact fate determined probabilistically by the fitted model). Both the population and spatial transition dynamics were modeled for all seven years of the study, with these processes only being observed in years 1 (2009), 4 (2012), and 7 (2015).

*Observation model*—We assumed the number of counts of each individual (*i*) in each trap (*j*) in the single sampling occasion per year (*t*) was a sex-specific, decreasing function of the distance between an individual’s yearly activity center, *s*_*it*_, and a trap location *x_j_*: 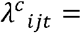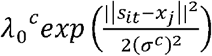, for *c* ∈ (*m,f*). Here, *λ*_0_^*m*^, 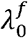, *σ*^*m*^, and *σ*^*f*^ are the male and female-specific baseline detection rate and spatial scale parameters, respectively. Further, we assumed theses counts were Poisson random variables. To limit confusion, we emphasize here that we need three instances of Greek lambda: *λ*_0_ for the baseline detection rate and *λ*^*c*^_*ijt*_ for the individual by trap by year detection rate, and λ for the population growth rate (described in Supplement B).

*Inference via data augmentation and MCMC*—Following Gardner et al. (2010), we augmented the observed data up to dimension *M*=800, chosen to be much greater than the total number of individuals ever alive over the 7 years, with *M* possible individuals distributed with equal density across patches so that that each patch had the same expected population density as the overall expected population density. We then introduce the *M* × *T* matrix ***z***, with entries 1 if individual *i* is alive and in the population in year *t* and 0 otherwise. This **z** matrix was used to estimate both the population size in all years and the population dynamics parameters as described by Chandler & Clark (2014) except that the sexes of the augmented and sex-indeterminate individuals needed to be updated on each MCMC iteration. This was accomplished by also augmenting the vector ***C*** up to dimension *M*, introducing a parameter for the probability that an individual is female, *p*^*sex*^, and assuming *c*_*i*_~*Bernoulli*(*p*^*sex*^). Then, using custom MCMC algorithms from the *OpenPopSCR* **R** package, we sampled from the joint posterior using 32 chains for 3000 iterations each, discarding 500 iterations as burn in for each chain. The remaining 64,000 samples were pooled for posterior inference. We used the typical vague priors for all model parameters (Chandler & Clark 2014, Royle et al. 2014) and the prior of uniform density across the state space implied that the priors for patch-level abundance, density, and sex ratio was uniform across patches. Posterior modes and 95% highest posterior density (HPD) intervals are reported throughout. A description of the derived parameter estimates, including population growth rates, patch-level quantities, and a post-hoc movement distance characterization can be found in Supplement B. The MCMC algorithms used can be found in the Data Supplement, along with the capercaillie data set.

## Results

In the years 2009, 2012 and 2015, a total of 259 capercaillies were detected at least once: 143 males, 112 females, and 4 individuals of unknown sex, with the percentage of detected individuals that were female in each year being 37, 38, and 46% (Table 1). The ratio of the observed number of individuals (Table 1) to the estimates of realized abundance (Supplement C; Table S3.1) suggests that between 90 and 92% of the total population was detected each year—95% of the male population, and between 80 and 88% of the female population. The overall detection probability varied by sex, with males estimated to be slightly more detectable than females (*λ*_0_^*m*^ = 1.193; 1.091 - 1.315 vs. *λ*_0_^*f*^ = 1.097; 0.940 - 1.303; Table 2). The estimate of σ for males was 1.44 times as large as that for females (Table 2), with a posterior probability that males had a larger detection spatial scale parameter of 1.

**Table 1.** Total and sex-specific number of individuals observed during each survey year.

| Year | Total | Male | Female | Unknown |
| --- | --- | --- | --- | --- |
| 2009 | 127 | 78 | 46 | 3 |
| 2012 | 111 | 69 | 42 | 0 |
| 2015 | 110 | 59 | 50 | 1 |

**Table 2.**
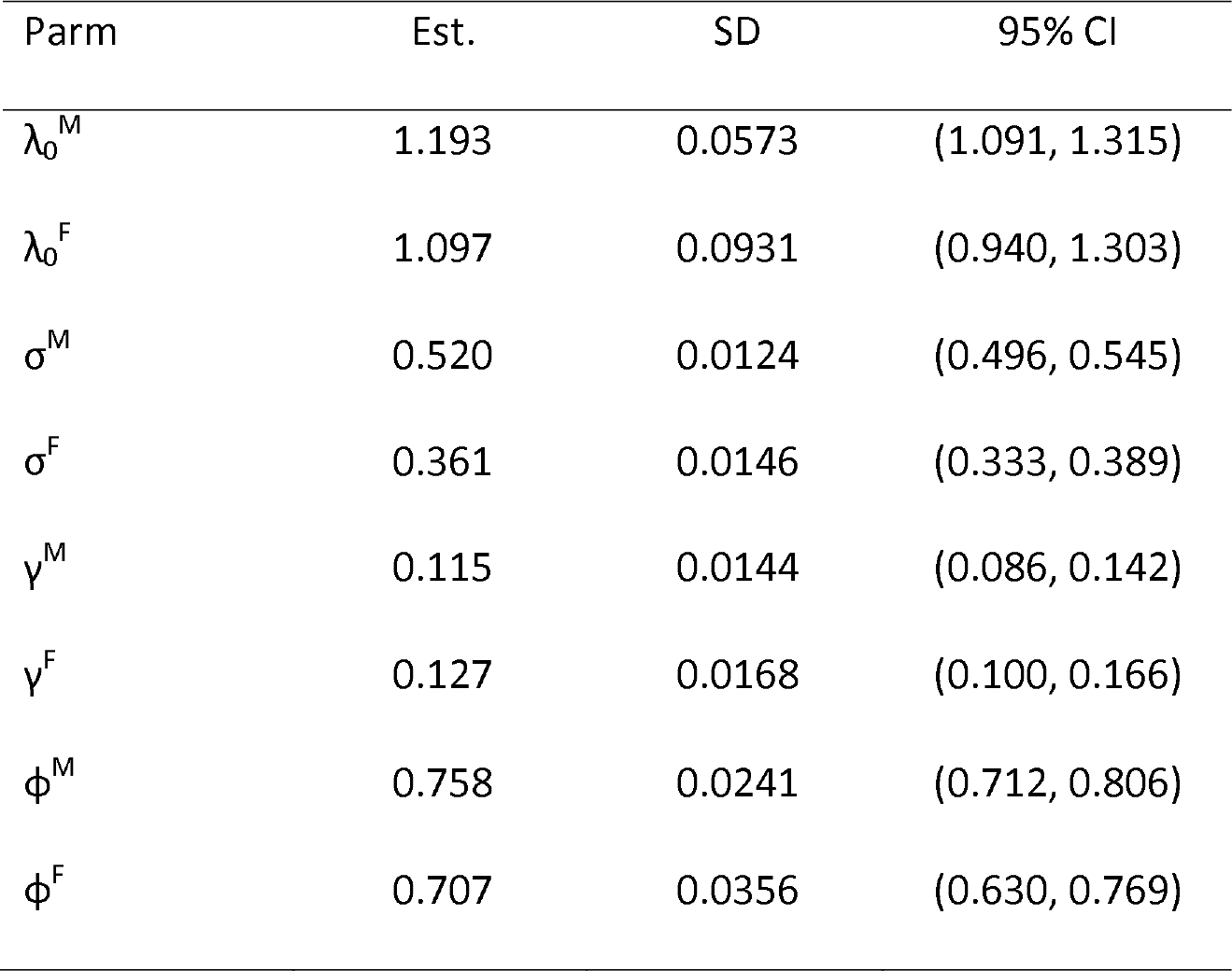
Posterior summaries for the detection and population dynamics parameters in the open-population SCR model with independent activity centers fit to Swiss capercaillie data over 7 years (2009-2015, with data observed in 2009, 2012, 2015). We show the posterior mode as a point estimate and the posterior standard deviation as an analogue to the standard error and, in the final two columns, the lower and upper credible interval limits (LCL and UCL, respectively) from a 95% highest posterior density interval. λ_0_ is the baseline detection rate, σ is the detection function spatial scale parameter, γ is per capita recruitment rate, and ϕ is yearly survival probability. Uppercase letters M and F denote estimates for males and females, respectively.

Annual per-capita recruitment rates were estimated to be 10% larger for females than males (Table 2), with a posterior probability of 0.8 that female per capita recruitment was larger. Annual survival probability was estimated to be larger for males than females (Table 2) with a posterior probability of 0.92. The yearly expected population growth rates, E[λ_t_], implied by these per capita recruitment and survival estimates are shown in Figure 3 and the associated expected abundances in the observed years and predicted into the future are shown in Figure 4. While the point estimates for population growth rate (Fig 3) and expected abundance (Fig 4) indicated a decline, the interval estimates were consistent with a moderate decline to a moderate increase, precluding our ability to diagnose the population trajectory with confidence from these quantities.

**Figure 3:**
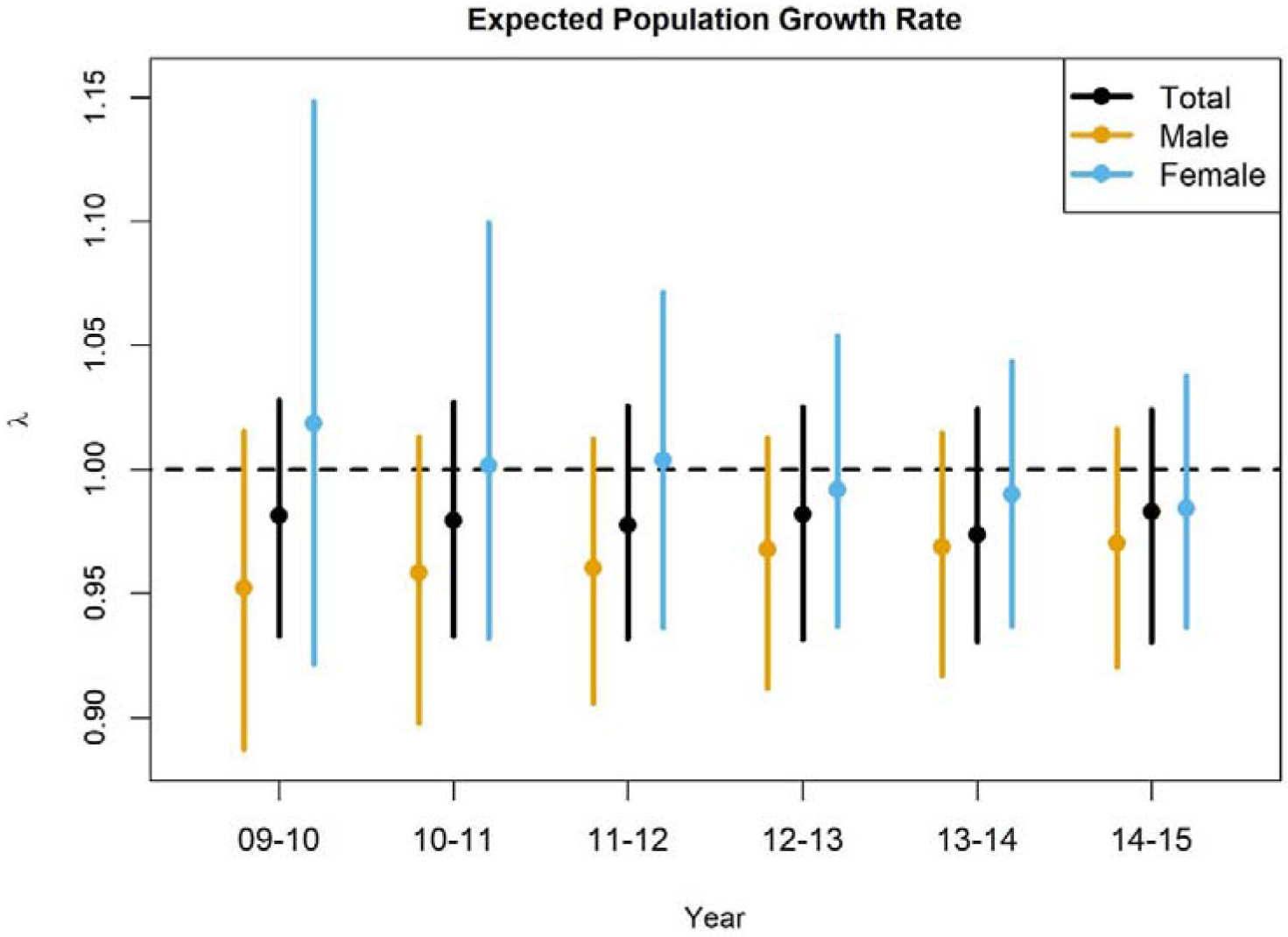
Total, male, and female expected population growth rate estimates between 2009 and 2015. Points indicate point estimates, and lines indicate 95% highest posterior density intervals.

**Figure 4:**
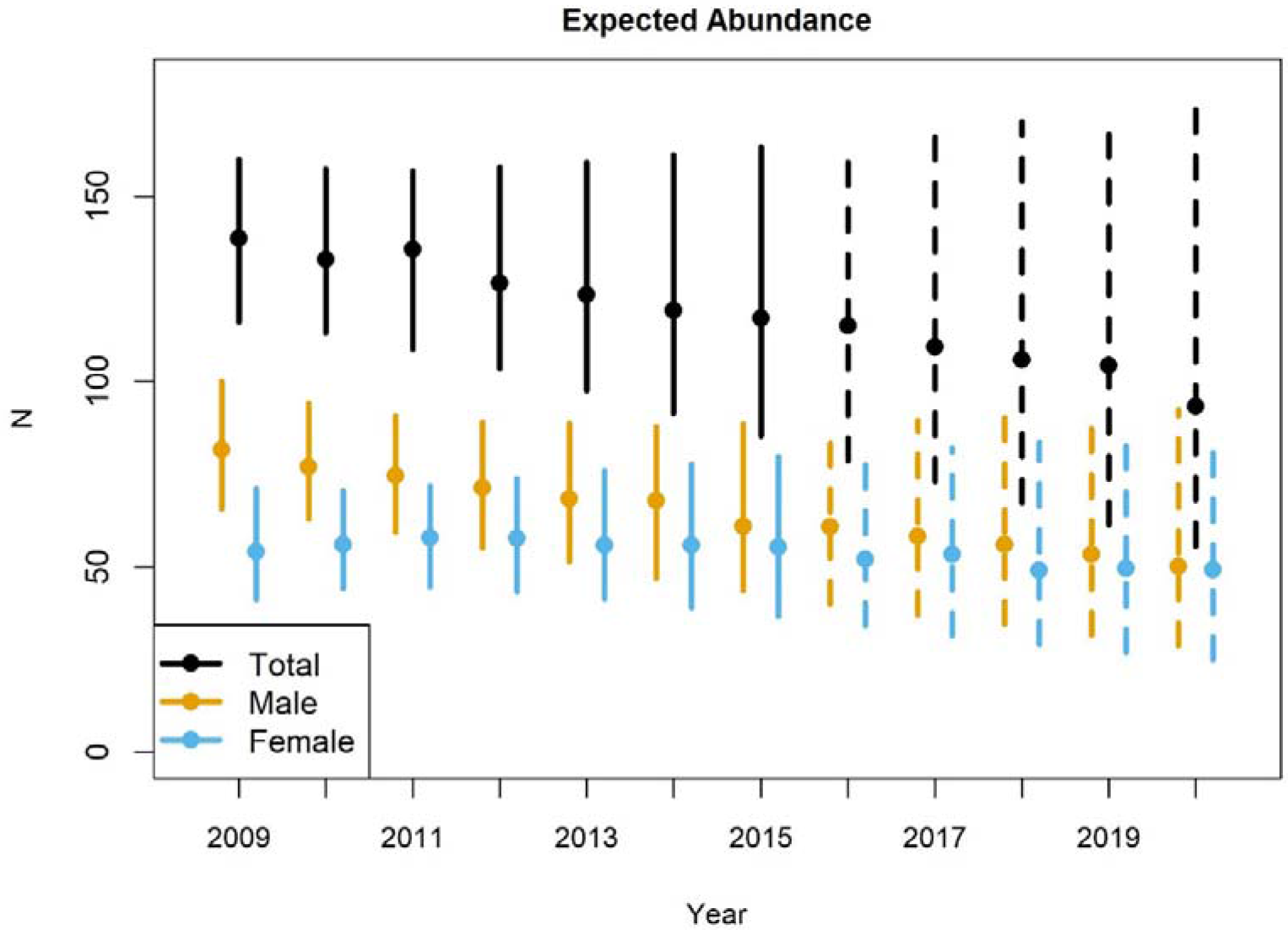
Total, male, and female expected abundance estimates between 2009 and 2015 and projected expected abundance estimates from 2016 to 2020. Points indicate point estimates, and lines indicate 95% highest posterior density intervals. Years concurrent with the collected data sets are indicated with solid lines and those that are projections beyond the data are indicated with dotted lines.

The derived parameters pooling population growth rate information across the 7-year period (described in detail in Supplement B) were more precise. The realized population growth rates *over the entire 7 year period*, *λ*_*T*_, were 0.88 (0.82 - 0.94), 0.76 (0.71 - 0.82), and 0.99 (0.91 - 1.21) for all individuals, males, and females, respectively. The *annual* realized population growth rates implied from the *λ*_*T*_ estimates assuming constant growth through time, *λ*_*fixed*_, were 0.98 (0.97 - 0.99), 0.95 (0.94 - 0.97), and 1.00 (0.99 - 1.03) for all individuals, males, and females, respectively. Using the realized population growth rates over the entire 7-year period, *λ*_*T*_, the posterior probability that total population, males, and females declined was 1.00, 1.00, and 0.20, respectively. The trend in realized density point estimates largely matched that seen in the expected abundance (Fig. 5; see Table S3.1 in Supplement C for realized abundance estimates).

**Figure 5:**
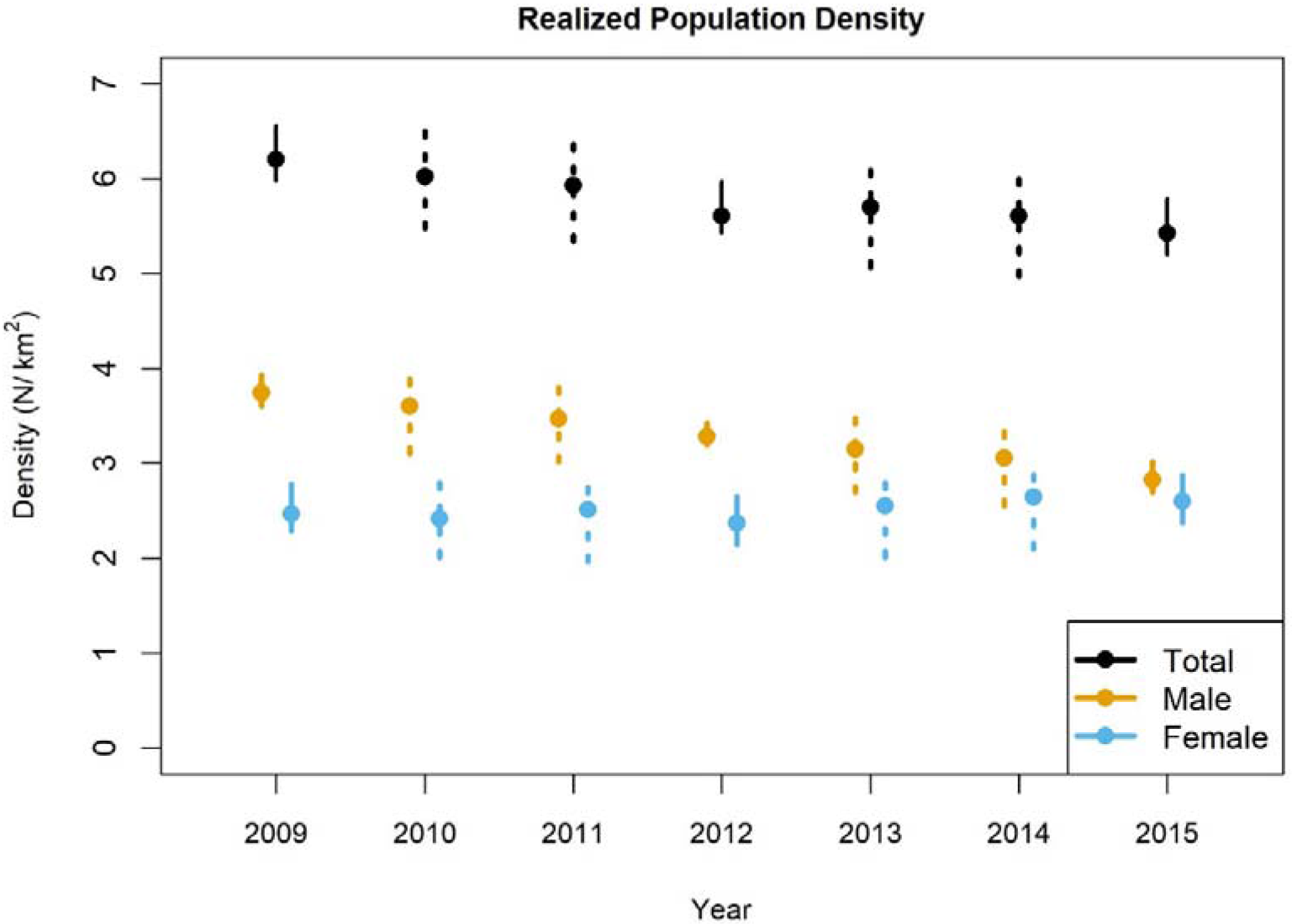
Total, male and female realized population density estimates between 2009 and 2015. Points indicate point estimates and lines indicate 95% highest posterior density intervals. Solid lines indicate years with data collection, dotted lines years without data collection.

Nearly half of the population was estimated to live in one of the eight patches (patch 4; Figure S3, Supplement C), with the majority of the population estimated to live in patches 4 and 6. The estimated percentage of the population living in these two patches declined from 71% in 2009 to 63% in 2015. Five of the eight patches consistently had an estimated realized density greater than 4 individuals per km^2^ (patches 4-8; Table S3.2, Supplement C). Densities in these higher density patches were estimated to have declined over the 7-year period in three patches (4, 6, and 8), increased in 1 patch (7), and remained roughly constant in patch 5. Total abundance mainly declined due to a decrease in the largest, most central patch (4) and male abundance primarily declined in the two largest patches, 4 and 6. Among the 5 higher density patches, two started in 2009 with a female-biased sex ratio (patches 5 and 8) and two others with male-biased sex ratio (patches 4 and 6). Sex ratios in the 5 higher density patches changed through time in patches 6 – 8, with patch 6 transitioning from male to female-biased.

We observed 85 potential movement events (individuals captured in consecutive 3-year periods) from 69 individuals. The mean 3-year movement for these 85 events was 1.02 km, with the vast majority of these estimated movement distances were less than 3 km (Figure 6), which is small compared to the distances between patches (mean minimum neighbour distance = 11 km, only 2 MMND < 3km). Thus, only 3 estimated movement distances (2 male, 1 female) were large enough to allow for movement between patches and only 2 (1 male, 1 female) were actually observed to move between patches. Given that we estimated that almost all of the individuals were captured, it is unlikely that we missed any long distance patch transitions.

**Figure 6:**
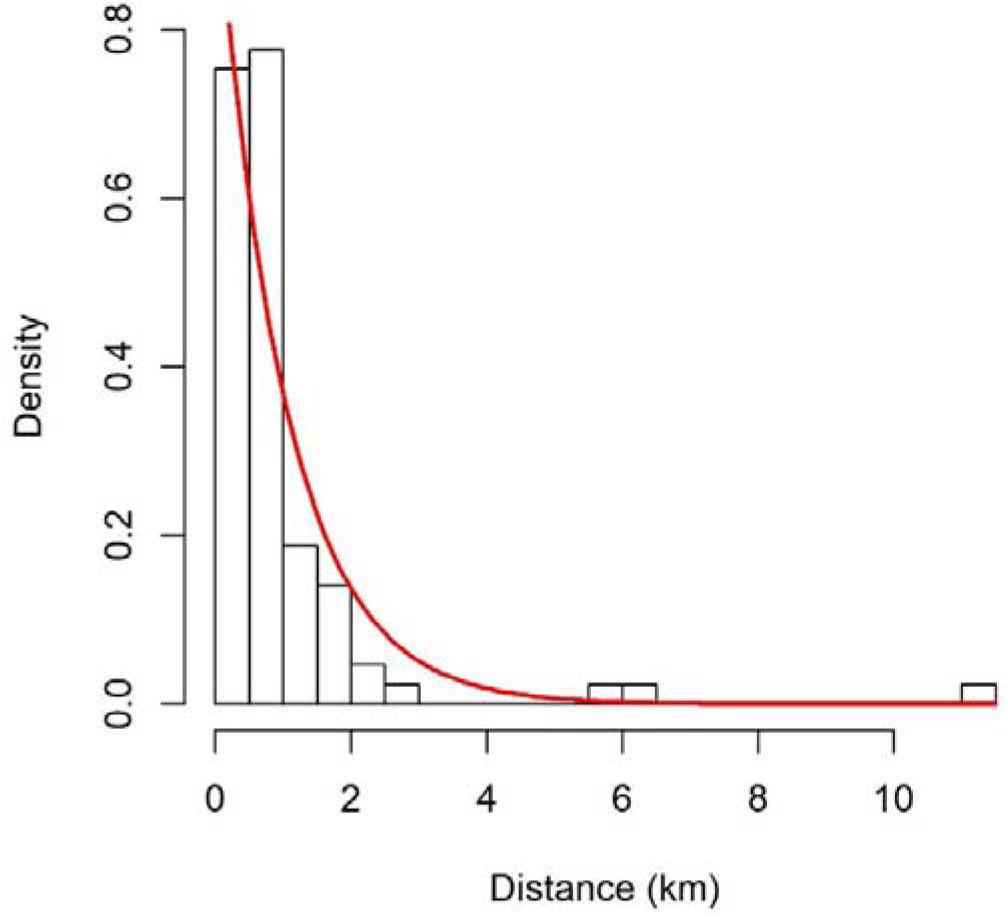
Estimated three year movement distances for 85 movement events for individuals detected in consecutive data sets (i.e. detected in 2009 and 2012 and/or in 2012 and 2015). Red line is a fitted exponential distribution 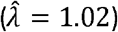

## Discussion

We applied a fully sex-specific open population SCR model that allows for population redistribution across a patchy habitat using a discrete state space to produce the first estimates of survival, recruitment, and population growth rates for the endangered capercaillie. Our approach yielded patch-level inferences about abundance, density, and sex-specific population dynamics, showing that 1) the observed population decline was attributed to a declining trend in males, with females, typically the more important sex for population persistence, remaining stable, and 2) the observed decline in the male population was attributed to only two of the eight habitat patches. The use of a hierarchical Bayesian open population model estimated using data augmentation allowed us to estimate annual survival, per capita recruitment, population growth rates, and abundance when surveys did not take place in every year. Further, the use of data augmentation allowed us to produce the patch by sex by year realized quantities (e.g., abundance, sex ratio) without the need to specify a model for the expected patch-level dynamics explicitly and estimate its parameters with sparse patch-level data.

Our assumption that individuals may relocate their activity centers independently between years within the patch they live, which we used to simplify the model for population redistribution across the patchy habitat, has an implication worth discussing. This activity center model removes a desirable feature of open population SCR models with explicit movement models—that exposure to capture in year *t+1* depends on the individual activity center location in year *t.* Removing this dependence erodes the ability to separate movement from survival and recruitment. However, Gardner et al. (2018) demonstrate that survival (and by implication, recruitment) can be estimated without bias using an independent activity center model when the state space extent is known and sampled exhaustively. We believe our inferences under the patch-constrained independent activity center model should be reliable for this data set because 1) we estimated that we detected >90% of all individuals in each year surveyed, 2) the scat survey covered the entire state space—the area in which cappercaillie were documented to occur between 1978 and 2009, and 3) we observed evidence of very little movement between patches that would allow individuals to enter and leave this set of 8 patches (discussed further below). Thus, we believe our treatment of individual activity centers was appropriate, but may be of limited use for other populations that are less thoroughly surveyed and for which a smaller proportion of individuals are captured in each year. The two advantages of constraining the independent activity centers to single patches instead of allowing them to relocate independently across the entire state space was 1) a more realistic quantification of exposure to capture for individuals that were not captured in the patch they live in a particular year and 2) a more realistic quantification of patch-level (meta)population dynamics.

We found moderate to strong evidence for sex-specific population dynamics—survival was estimated to be higher in males than females (posterior probability = 0.92) and per capita recruitment was estimated to be higher in females than males (0.80). These differences in demographic rates suggest that while the expected yearly population growth rate, E[λ_l_], for the sexes combined remained stable at around 0.98, for females the rate declined from around 1.02 to 0.98, while for the males the rate increased from 0.95 to 0.96 (Fig. 3). Corresponding to this pattern, the estimated total population size declined during the 7 years of the study and is projected to continue to decline. This pattern was driven by the male component of the population, with the female component of the population having remained roughly stable during the 7 years of the study and projected to remain stable over the next 5 years.

One counterintuitive result of this sex-specific population growth rate model is that males were estimated to have a higher survival probability and a lower per capita recruitment rate than females, with the former being a larger absolute difference. This appears to be in conflict with a declining male population and a stable female population; however, this perception is largely a consequence of the magnitude of male-bias in abundance in the first year of the survey. If we begin with the estimated expected abundances of 139, 82, and 54 for total, males, and females, respectively, we expect the number of males in the next year to be 82 × 0.756 + 139 × 0.115 = 61.992 + 15.985 = 77.977 and the expected number of females to be 54 × 0.707 + 139 × 0.127 = 38.178 + 17.653 = 55.831 (following the process model equation found in the Methods). The males lost 20.0 individuals due to death and gained 16.0 from recruitment, while the females only lost 15.8 to death and gained 17.7. Therefore, the greater number of males exposed to mortality led to more males dying than females and the lower male per capita recruitment rate ensured they could not make up the difference via recruitment. As the expected sex ratio converges toward 50:50 through time (Fig. 4, male vs. female N) and the differences in sex-specific abundance decrease, the differences in the sex-specific expected population growth rates become less extreme (Fig. 3) because the difference in sex-specific deaths decreases.

Despite the trends in the expected population growth rates and expected abundances, there was considerable uncertainty in the interval estimates (Fig. 3 and Fig. 4), so we could not rule out the equality of sex-specific population trajectories, or discriminate between the hypotheses of an increasing, stable, or declining population for either sex using these metrics. Therefore, we used the yearly realized population growth rate, *λ*_*fixed*_, calculated using the realized abundance in the first and last year of the survey, assuming constant population growth through time (see Supplement B for full derivation), as our best estimate of the population trajectory over the 7 year period. The point estimates for *λ*_*fixed*_, were 0.98, 0.95, and 1.00 for total population, males, and females, respectively. The posterior probabilities of each of these population growth rates being <1 were 1.0, 1.0, and 0.2, and the posterior probabilities that these population growth rates were <0.95 were 0.0, 0.25, and 0.00. Therefore, we found strong evidence that the total and male populations declined, but not more than 5% per year, weak evidence that the female population declined, and no evidence that female population declined by more than 5% per year. Thus, our best population-level interpretation of the data is that the sex ratio in this population was male-biased in the first year of the survey, male abundance declined, and female abundance remained stable, which is consistent with the more imprecise and ambiguous yearly-level estimates described above.

At the patch level, the population-level trend can largely be attributed to the population dynamics in the two largest patches 4 and 6 (Fig. 2). Over 60% of the population lived in these two patches, with over 40% living in patch 4, and both patches 4 and 6 began with male-biased sex ratios. The loss in total abundance over the 7 year period was largely driven by a reduction in abundance in patch 4, and a disproportionate loss of males over females in patch 6 exacerbated the observed male-biased population decline. These male losses led to a female-biased sex ratio in patch 6, while the sex ratio in patch 4 remained male-biased, but to a lesser degree than at the beginning of the study. Declines of males and a shifting sex ratio from male to female-biased in patch 3 also contributed to the overall population trend, but to a lesser degree than patches 4 and 6 due to a lower overall abundance. In sum, only 3 patches contributed to the overall population dynamics we observed. These patches are the largest, contiguous patches and the most centrally located among the metapopulation of patches (Fig. 2).

We used the spatial information contained in the capture locations to provide strong evidence that these 8 patches are spatially isolated. Of the 85 3-year activity center relocations, only 2 represented a transition between patches and only 3 relocated a distance that was greater than the minimum distance between patches. No observed 3-year relocations were large enough to allow for movement to or from patches in neighbouring populations outside of our study area. The estimated mean relocation distance over 3 years was 1 km, indicating that individuals relocate their activity centers very little, on average, between years. What these data cannot tell us about, however, is the extent of natal dispersal between patches because the surveys were conducted before birth of chicks.

While the main purpose of estimating population dynamics of this population was to assess the population status and trajectory of the endangered capercaillie, the parameter estimates can be used to better inform how this population and others like it are surveyed in the future. Mollet et al. (2015) investigated the impact of reducing the sampling intensity of this population on the estimates of population size and sex ratio in a single year, concluding that sampling effort could be reduced 25% while still yielding sufficient precision. With population dynamic parameter estimates now available, multi-year study designs with different inferential goals, such as assessing population growth rates can be compared via simulation. For example, designs may alter the number of years between surveys, survey different sets of patches in different years, or vary sampling intensity by patch. Other model elaborations may favour having some consecutive years surveyed, such as using Markov yearly activity center transitions or sex by year population dynamics parameters. Finally, the possibility exists of using a combination of scats with individual identities in some years and unidentified scat counts in others, similar to Chandler and Clark (2014); however, we suspect the density in this population is too large for unidentified counts to carry much information.

In closing, we utilized several recent developments in spatial and non-spatial open population models to analyse a challenging data set for the capercaillie to produce the first estimates of survival, recruitment, and population trajectory between 2009 and 2015, and producing population projections through 2020. Despite recent capercaillie population declines in Switzerland, the more important female component of the population appears stable. We recommend continued monitoring of this population at the current 3 year intervals, or with an adjusted study design which can be selected using the parameter estimates provided in this manuscript.

## Acknowledgements

We thank the gamekeepers, foresters and hunters of the cantons Schwyz, Zug and Glarus for assistance in the field, and the offices for hunting and for forest of the canton Schwyz for their help in organizing the field work.

## Author contributions

BA developed the open population models, MCMC algorithms, and software used in the analysis, led the data analysis, and contributed to writing of the manuscript. MK and CS contributed to model development, data analysis, and writing the manuscript. JOM and GP analyzed the genetic data and contributed to writing the manuscript, PM led the field work, organized the work of the genetic laboratories and contributed to writing the manuscript.

## Data Accessibility

All data and code required to run the analyses can be found in the Data Supplement.

## Supplement A. DNA extraction, genotyping and population genetic analyses

### DNA extraction and genotyping

Extraction of DNA from and genotyping of scat samples in years 2009 and 2015 were done by Ecogenics GmbH in Switzerland, following the method described by Jacob et al. (2010) and Mollet et al. (2015). In short, 12 nuclear microsatellites were amplified in four multiplex PCRs and sex was determined by amplification of a sex-specific marker. In 2012, DNA extraction and genotyping were done by the Nevada Genomics Center, USA, with the 12 microsatellite markers and the sex-specific marker amplified in two multiplex PCRs. In 2012 two of the markers (sTuD7 and sTuD4) were not possible to score, leaving 10 markers for further analyses. In all years, amplifications of the same sample were repeated at least three times in order to account for genotyping errors that usually occur with non-invasive samples (Bonin et al. 2004). The results from these replicates were used to construct the consensus genotype of each sample: only concordant results among three replicates of a locus (or among two concordant replicates in case a third one was missing) were accepted. Loci that could not be scored more than twice and loci with discordant results among replicates were scored as missing. Samples with less than eight successfully genotyped loci and those assigned to other grouse species were excluded from the analysis. Results of the replicate amplifications of the same sample were also used to calculate genotyping errors for each sampling year. The list of consensus genotypes per sampled year was used to identify unique individuals, recaptures, and their probability of identity (PI and Pl_sib_, Waits et al. 2001) with the programs Cervus 3.0.3 (Kalinowski et al. 2007 for data of years 2009 and 2015) or the R-package allele match (Galpern et al. 2012 for data of year 2012). Multi-locus genotypes were considered to be identical if they shared all the alleles at all the loci, excluding loci with missing values. An allelic combination representing one or several identical genotypes was considered to be unique if it differed from all the other allelic combinations by at least two alleles (excluding missing values). Loci were checked for departure of Hardy-Weinberg equilibrium, null allele frequencies and measures of genetic diversity (number of alleles per locus, allelic richness) using GENEPOP on the web (http://genepop.curtin.edu.au; Raymond & Rousset 1995; Rousset 2008), Cervus 3.0.3 and FSTAT 2.9.3.2 (Goudet 2002).

The list of unique multi-locus genotypes of all surveyed years was used to identify recaptures during the study period using allele match. Identical genotypes at all 10 loci (N=63), mismatches due to 1 (N=35) or 2 (N=11) missing loci, or mismatches at 1 allele (N=4) were considered to belong to the same individual.

After these analyses, for each study year, we had a list of detections of individually identifiable birds with the following associated information: date, sex, and location, the latter also containing the specific patch in which an individual was located.

### Results of population genetic analyses

The basic measures of genetic diversity and Hardy-Weinberg equilibrium were similar across years (see Supplementary Table 1). Genotyping error rates differed among the single loci, with values ranging from 1% - 28%. Taking all loci together, the mean genotyping error rates were consistently low (10%, 2% and 6% for years 2009, 2012 and 2015, respectively, see Supplementary Table 2) across years. The proportion of incomplete genotypes varied among loci (from 5% - 60%) and years (7%, 33% and 8% for years 2009, 2012 and 2015, respectively, see Supplementary Table 2); although the year 2012 had a relatively high proportion of incomplete genotypes, it is important to note that more than half of these cases were due to one (out of three) unsuccessful PCR replicate, with the other two replicates showing concordant results.

**Supplementary Table S1:** Probability of identity (PI) and measures of genetic diversity at 10 microsatellite loci in our capercaillie population for the years 2009, 2012 and 2015 sorted by polymorphic information content (PIC).

| Year | Locus | N | A | R | H<sub>o</sub> | H<sub>e</sub> | PIC | PI | PI<sub>sib</sub> | HW | F<sub>Null</sub> |
| --- | --- | --- | --- | --- | --- | --- | --- | --- | --- | --- | --- |
| 2009 | sTuD5 | 129 | 9 | 8.977 | 0.837 | 0.856 | 0.835 | 0.039 | 0.334 | 0.783 | 0.009 |
|  | BG18 | 129 | 5 | 5 | 0.829 | 0.776 | 0.737 | 0.088 | 0.385 | 0.092 | -0.037 |
|  | sTuT4 | 129 | 5 | 5 | 0.674 | 0.761 | 0.718 | 0.098 | 0.395 | **0.065** | 0.057 |
|  | sTuD1 | 129 | 5 | 5 | 0.698 | 0.722 | 0.669 | 0.129 | 0.423 | 0.211 | 0.020 |
|  | sTuT2 | 129 | 4 | 4 | 0.535 | 0.571 | 0.524 | 0.230 | 0.523 | 0.292 | 0.029 |
|  | BG15 | 126 | 3 | 3 | 0.579 | 0.577 | 0.506 | 0.249 | 0.525 | **0.856** | 0.001 |
|  | sTuD6 | 128 | 10 | 9.999 | 0.547 | 0.503 | 0.485 | 0.265 | 0.566 | 0.984 | -0.065 |
|  | sTuT3 | 129 | 5 | 5 | 0.481 | 0.518 | 0.461 | 0.290 | 0.565 | 0.597 | 0.042 |
|  | sTuT1 | 128 | 4 | 3.984 | 0.344 | 0.497 | 0.445 | 0.305 | 0.579 | 0.000 | 0.169 |
|  | sTuD3 | 128 | 4 | 4 | 0.477 | 0.502 | 0.432 | 0.318 | 0.580 | 0.718 | 0.023 |
|  | Across loci<br>(SE) |  | 5.400<br>(0.718) | 5.396<br>(0.717) | 0.600<br>(0.05) | 0.628<br>(0.043) | 0.581<br>(0.046) | 1.85E-08 | 6.34E-04 |  |  |
| 2012 | sTuD5 | 99 | 11 | 11 | 0.818 | 0.860 | 0.839 | 0.037 | 0.331 | 0.377 | 0.022 |
|  | BG18 | 112 | 5 | 5 | 0.714 | 0.765 | 0.722 | 0.096 | 0.393 | 0.001 | 0.032 |
|  | sTuT4 | 124 | 6 | 5.960 | 0.742 | 0.756 | 0.712 | 0.102 | 0.399 | 0.096 | 0.005 |
|  | sTuD1 | 127 | 6 | 5.952 | 0.677 | 0.736 | 0.686 | 0.118 | 0.413 | **0.171** | 0.039 |
|  | sTuT2 | 120 | 8 | 7.3 | 0.608 | 0.590 | 0.549 | 0.209 | 0.508 | **0.037** | -0.019 |
|  | sTuD6 | 118 | 11 | 10.649 | 0.619 | 0.561 | 0.541 | 0.212 | 0.524 | 0.867 | -0.072 |
|  | BG15 | 125 | 5 | 4.584 | 0.616 | 0.575 | 0.506 | 0.249 | 0.526 | 0.019 | -0.039 |
|  | sTuT1 | 111 | 6 | 5.784 | 0.360 | 0.537 | 0.487 | 0.264 | 0.549 | 0.000 | 0.182 |
|  | sTuD3 | 124 | 4 | 3.798 | 0.565 | 0.509 | 0.429 | 0.321 | 0.577 | 0.344 | -0.064 |
|  | sTuT3 | 121 | 5 | 5 | 0.471 | 0.450 | 0.401 | 0.351 | 0.614 | 0.678 | -0.024 |
|  | Across loci |  | 6.700 | 6.503 | 0.619 | 0.634 | 0.587 | 1.40E- | 5.84E- |  |  |
|  | (SE) |  | (0.790) | (0.779) | (0.042) | (0.043) | (0.046) | 08 | 04 |  |  |
| 2015 | sTuD5 | 99 | 11 | 9.973 | 0.848 | 0.842 | 0.819 | 0.046 | 0.342 | 0.248 | -0.005 |
|  | BG18 | 112 | 5 | 5 | 0.761 | 0.766 | 0.723 | 0.095 | 0.393 | **0.174** | -0.004 |
|  | sTuT4 | 124 | 6 | 5 | 0.732 | 0.757 | 0.713 | 0.102 | 0.398 | **0.688** | 0.016 |
|  | sTuD1 | 127 | 6 | 5 | 0.782 | 0.725 | 0.672 | 0.127 | 0.421 | 0.070 | -0.041 |
|  | sTuD6 | 118 | 11 | 10 | 0.563 | 0.539 | 0.523 | 0.228 | 0.539 | 0.599 | -0.033 |
|  | sTuT2 | 120 | 8 | 4 | 0.563 | 0.564 | 0.514 | 0.239 | 0.529 | 0.028 | 0.007 |
|  | sTuT1 | 111 | 6 | 3.973 | 0.321 | 0.573 | 0.507 | 0.248 | 0.527 | 0.000 | 0.274 |
|  | BG15 | 125 | 5 | 3 | 0.640 | 0.555 | 0.492 | 0.260 | 0.539 | 0.113 | -0.084 |
|  | sTuT3 | 121 | 5 | 5 | 0.495 | 0.507 | 0.454 | 0.296 | 0.572 | 0.020 | 0.016 |
|  | sTuD3 | 124 | 4 | 3 | 0.536 | 0.478 | 0.392 | 0.359 | 0.602 | 0.438 | -0.062 |
|  | Across loci |  | 6.700 | 5.395 | 0.624 | 0.631 | 0.581 | 2.11E- | 6.28E- |  |  |
|  | (SE) |  | (0.79) | (0.804) | (0.050) | (0.041) | (0.044) | 08 | 04 |  |  |

**Supplementary Table S2:**
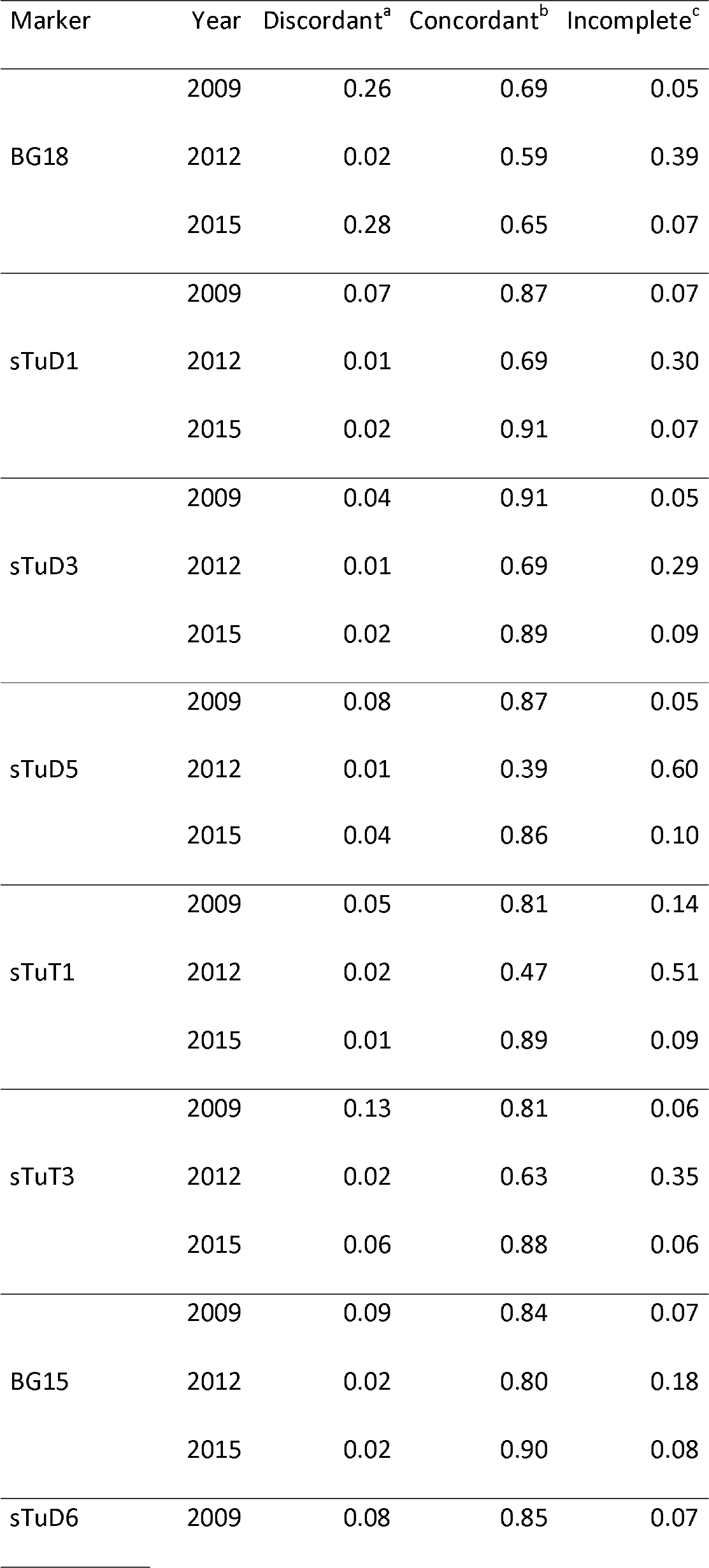
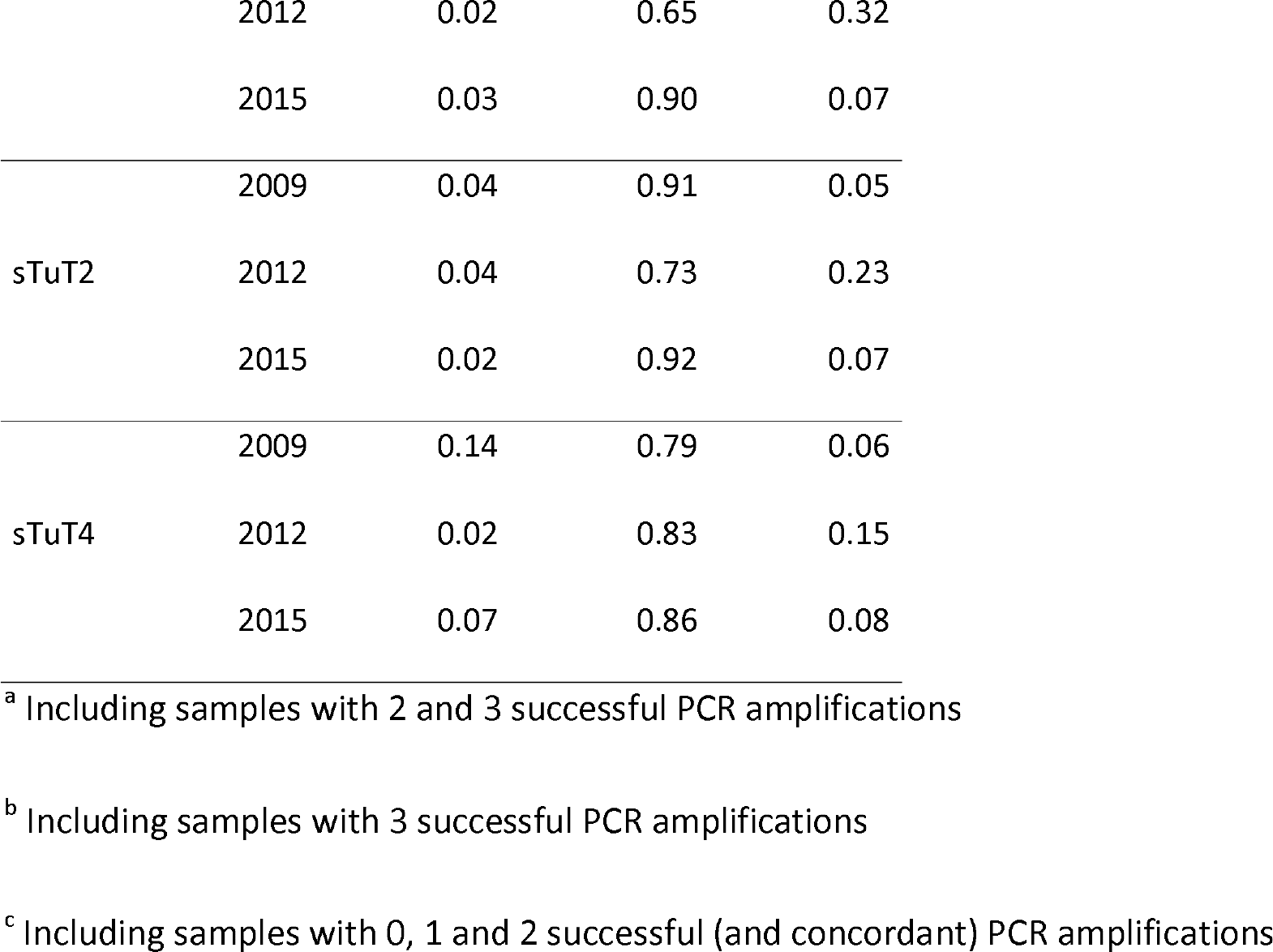
Comparison of genotypes obtained from three replicate PCRs of the same sample during the study years (N 2009 = 587, N 2012 = 628, N 2015 = 561). The proportion of each outcome is shown.

## Supplement B. Derived parameters in Spatial Capture-recapture

We refer readers to Chandler and Clark (2014) for the full details of how the ***z*** matrix is used for inference about survival, recruitment, and abundance parameters. Here, we will provide some more detail to describe how various derived quantities were calculated. Both expected and realized abundance, density, and sex ratio in hierarchical open population models can be estimated. The expected quantities are often given directly by parameters in the model and reflect what *would* happen on average over many realizations of the population dynamics process under study. In contrast, the realized quantities are based on the estimates of the z matrix and reflect our best estimate of what *actually occurred* in the study population during the sampled period. We regard the *realized* quantities as of more interest to describe what population dynamics occurred while we were observing this population. However, we are also interested in the *expected* quantities for inferences about population trends through time and especially for projecting population trends into the future.

On each MCMC iteration, realized abundance in year *t* is obtained by summing the ***z*** subvector ***z***_t_ and the realized sex-specific abundances in year *t* are obtained by summing the elements of the subvector ***z***_t_ whose corresponding elements of the vector ***C*** are of the focal sex. More specifically, *N*_*t*_ = ∑_*i*_*z*_*it*_, *N*^*m*^_*t*_ = ∑_*i*_*z*_*it*_*I*(*C*_*i*_ = 0) and *N*^*f*^_*t*_ = ∑_*i*_*z*_*it*_*I*(*C*_*i*_ = 1), where I() is an indicator function taking value 1 if the interior condition is met and 0 otherwise. All of these realized abundances can be obtained by dividing the realized abundances by the area of the state space. The realized sex ratio (probability an individual is female) in year *t* is then 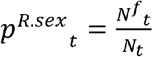 and the realized population growth rate estimates between year *t* and *t+1* are 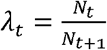, with the sex-specific realized population growth rates obtained by swapping in the sex-specific realized abundance estimates. Finally, patch and year-specific realized total and sex-specific abundance, density, and realized sex ratio estimates can be derived by using the **C** posteriors to determine sex, the ***s*** posteriors to determine the patch number each ***s***_it_ is located in, and the ***z*** posteriors to determine alive status.

Similarly, on each MCMC iteration, the expected versions of the quantities described above can be calculated. As in typical Bayesian SCR (Royle et al. 2014), the expected population size in year 1 is *E*[*N*_1_] = *ψM*, where *ψ* is the probability z_i_ is included in the population on occasion 1, assumed to follow *z*_*i*1_ = *Bernoulli*(*ψ*). The expected sex-specific population sizes in year 1 are then *E*[*N*^*f*^_1_] = *ψp^sex^M* and *E*[*N*^*m*^_1_] *ψ*(1 − *p*^*sex*^)*M* and for females and males, respectively. The expected sex-specific population size in years *2,…, L* are obtained by plugging the sex-specific expected abundances into the population growth difference equation in the main text and the expected total population size is just the sum of the expected sex-specific population sizes. Finally, the expected sex ratios, *p*^*E.sex*^_*t*_ and population growth rates, *E*[*λ*_*t*_], can be obtained using the same equations as used for the realized estimates, but plugging in the expected abundances.

We will report the realized quantities for sex and sex by patch density and sex ratio estimates as these best reflect the population dynamics that occurred during the survey. We report the expected population growth rates due to the fact that the realized population growth rates involve the estimates of the realized abundance estimates in the years not observed (2,3,4,6) and because the expected population growth rates best describe how population growth is estimated to change over the course of the 7 year duration of the survey. As our best estimate of the realized population growth rate over the entire survey, we calculated 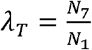, and under the assumption of a fixed realized population growth rate through time, we calculated the yearly mean realized population growth rate following 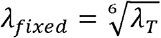, with the sixth root stemming from the fact that there are 6 abundance changes over 7 years. Finally, we use the expected abundance estimates to predict the population sizes 5 years into the future from the last year of the survey.

In order to support our assumption that no individuals moved between patches except for the two that were observed to transition between patches once each, we calculated the posterior modes of the “observed” dispersal distances (3-year dispersal distances for individuals observed in consecutive surveys) by post-processing the ***s*** and ***z*** posteriors for these individuals. We then estimated the rate parameter of an exponential dispersal kernel using the posterior modes of the dispersal distances via maximum likelihood. Because these dispersal distances are over a 3 year period, we assume they are larger than the yearly dispersal distances and so our assumption of no movement between patches will be supported if we observe very few estimated dispersal distances larger than the minimum distance between patches.

## Supplement C. Sex and sex by patch level abundance and density estimates

**Table S3.1.** Posterior summaries for the realized abundance parameters (Est.) in the open-population SCR model with independent activity centers fit to Swiss capercaillie data over 7 years (2009-2015, with data observed in 2009, 2012, 2015). We show the posterior mode as a point estimate and the posterior standard deviation as an analogue to the standard error and, in the final two columns, the lower and upper credible interval limits (LCL and UCL, respectively) from a 95% highest posterior density interval. N_1-7_ is abundance in years 1 to 7. Uppercase letters M and F denote estimates for males and females, respectively.

| Parameter | Estimate | SD | 95% CI |
| --- | --- | --- | --- |
| $N_1$ | 136 | 3.48 | (131, 144) |
| $N_2$ | 132 | 6.78 | (120, 146) |
| $N_3$ | 130 | 6.63 | (117, 142) |
| $N_4$ | 123 | 3.25 | (119, 131) |
| $N_5$ | 125 | 6.55 | (111, 136) |
| $N_6$ | 123 | 6.53 | (109, 134) |
| $N_7$ | 119 | 3.46 | (114, 127) |
| $N_1^M$ | 82 | 2.06 | (79, 86) |
| $N_2^M$ | 79 | 4.90 | (68, 87) |
| $N_3^M$ | 76 | 4.73 | (66, 84) |
| $N_4^M$ | 72 | 1.59 | (70, 75) |
| $N_5^M$ | 69 | 4.69 | (59, 77) |
| $N_6^M$ | 67 | 4.64 | (56, 74) |
| $N_7^M$ | 62 | 1.83 | (59, 66) |
| $N_1^F$ | 54 | 3.00 | (50, 61) |
| $N_2^F$ | 53 | 5.23 | (44, 64) |
| $N_3^F$ | 55 | 5.14 | (43, 63) |
| $N_4^F$ | 52 | 2.86 | (47, 58) |
| $N_5^F$ | 56 | 5.16 | (44, 64) |
| $N_6^F$ | 58 | 5.17 | (46, 66) |
| $N_7^F$ | 57 | 2.99 | (52, 63) |

**Table S3.2.** Year- and patch-specific total (T), male (M) and female (F) realized densities, as well as realized sex ratio (probability female, SR) estimates. Point estimates are posterior modes and interval estimates are 95% highest posterior density (HPD) intervals. Trend indicates no change (=), an increase (+), or a decrease (−), with increases and decreases indicated when the posterior probabilities of the difference between years 2009 and 2015 were > 0.90 for either an increase or a decrease. Sex ratio estimates for which the 95% HPD intervals did not overlap 0.5 are in bold. Note, sex-specific abundance posterior modes do not add exactly to the total abundance posterior modes.

| Patch | Year |  |  |  |  |  |  | Trend |
| --- | --- | --- | --- | --- | --- | --- | --- | --- |
| – | 2009 | 2010 | 2011 | 2012 | 2013 | 2014 | 2015 | (prob) |
| Parm |  |  |  |  |  |  |  |  |
| 1-T | 0.00 (0.00 – 2.91) | 0.98 (0.00 – 3.88) | 0.97 (0.00 – 3.88) | 0.97 (0.97 – 2.91) | 1.94 (0.97 – 3.88) | 1.95 (0.97 – 3.88) | 0.97 (0.97 – 2.91) | = |
| 2-T | 1.74 (1.74 – 3.48) | 2.61 (0.87 – 5.22) | 3.48 (0.87 – 5.22) | 3.48 (3.48 – 4.35) | 3.48 (1.74 – 6.09) | 3.48 (1.74 – 6.09) | 3.48 (3.48 – 5.22) | + (0.96) |
| 3-T | 2.15 (2.15 – 2.77) | 3.08 (2.15 – 4.62) | 4.00 (2.77 – 5.23) | 4.62 (4.62 – 5.23) | 3.69 (2.46 – 5.23) | 3.08 (1.85 – 4.62) | 2.46 (2.46 – 3.08) | = |
| 4-T | 7.96 (7.71 – 8.33) | 7.22 (6.09 – 8.21) | 6.84 (5.85 – 7.84) | 6.10 (5.97 – 6.59) | 6.10 (5.22 – 7.09) | 6.10 (5.22 – 7.09) | 5.97 (5.85 – 6.34) | - (1.00) |
| 5-T | 4.63 (4.63 – 6.48) | 5.56 (3.70 – 8.33) | 5.56 (3.70 – 8.33) | 6.48 (6.48 – 8.33) | 6.47 (3.70 – 9.26) | 5.55 (2.78 – 8.33) | 4.63 (4.63 – 7.41) | = |
| 6-T | 7.79 (7.79 – 8.57) | 7.01 (5.71 – 8.31) | 5.98 (4.94 – 7.53) | 4.94 (4.68 – 5.45) | 5.20 (4.16 – 6.75) | 5.71 (4.42 – 7.27) | 6.49 (6.23 – 7.01) | - (1.00) |
| 7-T | 4.61 (4.15 – 5.53) | 4.61 (3.23 – 6.91) | 5.53 (3.23 – 6.91) | 5.53 (5.07 – 6.45) | 5.53 (3.23 – 7.37) | 5.99 (3.69 – 7.83) | 5.99 (5.53 – 6.91) | + (0.93) |
| 8-T | 9.63 (8.89 – 11.85) | 9.63 (5.93 – 12.59) | 8.89 (5.93 – 11.85) | 8.89 (8.15 – 10.37) | 8.15 (5.19 – 11.85) | 8.15 (4.44 – 11.11) | 8.15 (6.67 – 9.63) | - (0.92) |
| 1-M | 0.00 (0.00 – 0.97) | 0.00 (0.00 – 1.94) | 0.00 (0.00 – 1.94) | 0.00 (0.00 – 0.97) | 0.00 (0.00 – 1.94) | 0.00 (0.00 – 1.94) | 0.00 (0.00 – 0.97) | = |
| 2-M | 0.87 (0.87 – 1.74) | 1.74 (0.00 – 3.48) | 1.74 (0.00 – 3.48) | 2.61 (2.61 – 3.48) | 2.61 (1.74 – 4.35) | 2.61 (1.74 – 4.35) | 2.61 (2.61 – 3.48) | + (1.00) |
| 3-M | 1.23 (1.23 – 1.54) | 1.54 (0.92 – 2.46) | 1.85 (1.23 – 2.77) | 2.46 (2.46 – 2.77) | 1.85 (0.92 – 2.77) | 1.54 (0.62 – 2.46) | 0.92 (0.92 – 1.23) | - (0.89) |
| 4-M | 5.22 (5.10 – 5.47) | 4.73 (3.98 – 5.47) | 4.23 (3.61 – 5.10) | 4.10 (4.10 – 4.23) | 4.10 (3.36 – 4.60) | 3.98 (3.36 – 4.60) | 3.73 (3.73 – 3.86) | - (1.00) |
| 5-M | 1.86 (1.85 – 3.70) | 1.86 (0.93 – 3.70) | 2.78 (0.93 – 4.63) | 2.78 (2.78 – 3.70) | 2.78 (0.93 – 4.63) | 1.85 (0.93 – 4.63) | 1.85 (1.85 – 3.70) | = |
| 6-M | 5.71 (5.71 – 5.97) | 4.93 (3.90 – 5.97) | 3.64 (2.86 – 4.68) | 2.86 (2.86 – 3.13) | 2.85 (1.82 – 3.64) | 2.59 (1.56 – 3.38) | 2.34 (2.34 – 2.86) | - (1.00) |
| 7-M | 1.84 (1.84 – 2.76) | 2.30 (1.38 – 3.69) | 2.77 (1.84 – 4.15) | 3.22 (3.23 – 3.69) | 2.76 (1.84 – 5.07) | 3.22 (1.84 – 4.61) | 3.23 (3.23 – 4.15) | + (0.98) |
| 8-M | 2.96 (2.96 – 4.44) | 3.70 (2.22 – 5.93) | 3.70 (2.22 – 5.93) | 3.70 (3.70 – 5.19) | 3.71 (2.22 – 5.93) | 3.70 (2.22 – 5.93) | 3.70 (3.70 – 5.19) | = |
| 1-F | 0.00 (0.00 – 1.94) | 0.98 (0.00 – 2.91) | 0.97 (0.00 – 2.91) | 0.97 (0.00 – 1.94) | 0.97 (0.00 – 2.91) | 0.97 (0.00 – 2.91) | 0.97 (0.00 – 2.91) | = |
| 2-F | 0.87 (0.87 – 1.74) | 0.87 (0.87 – 2.61) | 0.87 (0.87 – 2.61) | 0.87 (0.87 – 1.74) | 0.87 (0.00 – 2.61) | 0.87 (0.00 – 2.61) | 0.87 (0.00 – 2.91) | = |
| 3-F | 0.92 (0.92 – 1.54) | 1.54 (0.62 – 2.46) | 1.85 (0.62 – 2.77) | 2.15 (2.15 – 2.77) | 2.16 (1.23 – 3.38) | 1.85 (0.62 – 2.77) | 1.54 (1.54 – 2.15) | + (0.94) |
| 4-F | 2.61 (2.36 – 2.99) | 2.36 (1.74 – 3.11) | 2.24 (1.62 – 2.99) | 1.99 (1.87 – 2.36) | 2.11 (1.49 – 2.86) | 2.12 (1.62 – 2.99) | 2.24 (2.11 – 2.61) | - (0.94) |
| 5-F | 2.78 (2.78 – 4.63) | 2.78 (1.85 – 4.63) | 2.78 (1.85 – 4.63) | 3.71 (3.70 – 5.56) | 3.70 (1.85 – 5.56) | 2.77 (1.85 – 5.56) | 2.77 (2.78 – 4.63) | = |
| 6-F | 2.08 (1.82 – 2.60) | 2.08 (1.30 – 3.12) | 2.08 (1.56 – 3.38) | 1.82 (1.82 – 2.60) | 2.60 (1.56 – 3.64) | 3.12 (2.08 – 4.16) | 3.90 (3.64 – 4.42) | + (1.00) |
| 7-F | 2.31 (2.30 – 3.23) | 2.30 (0.92 – 3.69) | 2.31 (0.92 – 3.69) | 1.84 (1.84 – 3.23) | 2.30 (1.38 – 3.69) | 2.76 (1.38 – 4.15) | 2.30 (2.30 – 3.23) | = |
| 8-F | 5.93 (5.93 – 8.15) | 5.92 (2.96 – 8.15) | 5.19 (2.96 – 7.41) | 4.44 (4.44 – 6.67) | 4.45 (1.48 – 6.67) | 3.70 (0.74 – 5.93) | 3.70 (2.96 – 5.19) | - (0.98) |
| 1-SR | 1.00 (0.00 – 1.00) | 1.00 (0.00 – 1.00) | 1.00 (0.00 – 1.00) | 1.00 (0.50 – 1.00) | 1.00 (0.50 – 1.00) | 1.00 (0.50 – 1.00) | 1.00 (0.50 – 1.00) | = |
| 2-SR | 0.50 (0.33 – 0.67) | 0.50 (0.25 – 1.00) | 0.50 (0.17 – 0.75) | 0.25 (0.20 – 0.40) | 0.25 (0.00 – 0.57) | 0.25 (0.00 – 0.57) | 0.25 (0.20 – 0.40) | - (0.93) |
| 3-SR | 0.43 (0.38 – 0.56) | 0.50 (0.29 – 0.70) | 0.50 (0.30 – 0.67) | 0.47 (0.47 – 0.52) | 0.50 (0.33 – 0.75) | 0.50 (0.33 – 0.75) | **0.63** (0.56 – 0.70) | + (0.99) |
| 4-SR | **0.33** (0.31 – 0.37) | **0.34** (0.26 – 0.42) | **0.33** (0.26 – 0.43) | **0.33** (0.31 – 0.37) | **0.33** (0.26 – 0.43) | **0.33** (0.27 – 0.44) | **0.37** (0.35 – 0.41) | + (0.95) |
| 5-SR | **0.60** (0.50 – 0.71) | 0.50 (0.38 – 0.83) | 0.50 (0.33 – 0.75) | 0.57 (0.50 – 0.67) | 0.50 (0.38 – 0.83) | 0.50 (0.33 – 0.83) | 0.60 (0.50 – 0.78) | = |
| 6-SR | **0.27** (0.23 – 0.31) | **0.33** (0.21 – 0.43) | **0.33** (0.25 – 0.48) | **0.38** (0.37 – 0.48) | 0.50 (0.35 – 0.64) | 0.50 (0.42 – 0.69) | **0.62** (0.58 – 0.65) | + (1.00) |
| 7-SR | 0.56 (0.45 – 0.63) | 0.50 (0.29 – 0.70) | 0.49 (0.22 – 0.63) | **0.36** (0.31 – 0.47) | 0.50 (0.27 – 0.64) | 0.50 (0.29 – 0.64) | 0.41 (0.36 – 0.50) | - (0.94) |
| 8-SR | **0.67** (0.56 – 0.71) | 0.50 (0.42 – 0.77) | 0.50 (0.40 – 0.75) | **0.58** (0.50 – 0.64) | 0.50 (0.27 – 0.71) | 0.50 (0.27 – 0.73) | 0.50 (0.36 – 0.57) | - (0.99) |

**Figure S3.**
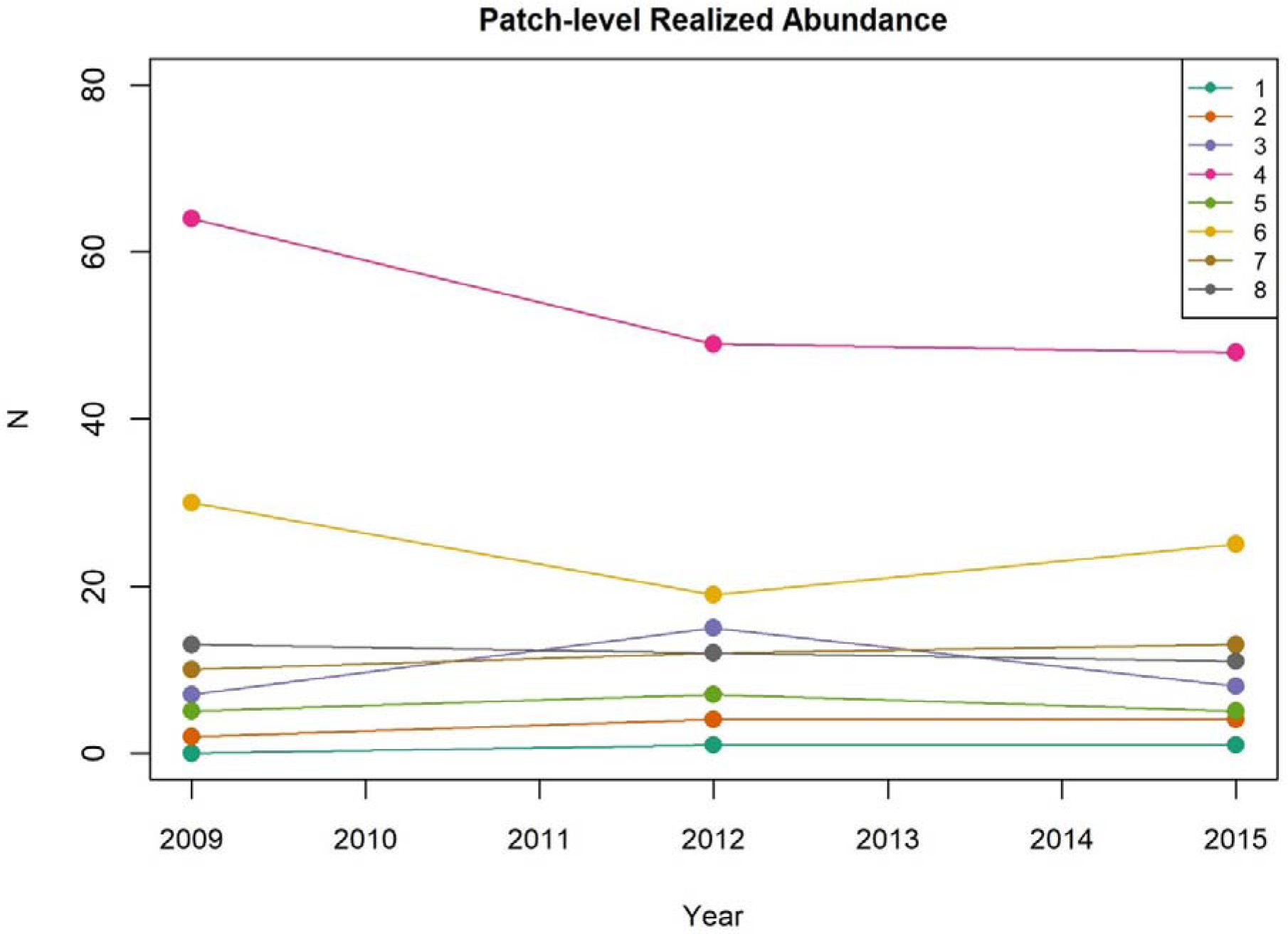
Patch (1-8) by year posterior modes of realized abundance.

## References

Augustine, B. C. (2018). OpenPopSCR, GitHub repository, https://github.com/benaug/OpenPopSCR.

Bollmann K, Weibel P, & Graf R. F. (2005), An analysis of central Alpine capercaillie spring habitat at the forest stand scale. Forest Ecology and Management, 215: 307–318. doi: 10.1016/j.foreco.2005.05.019

Borchers, D. L., & Efford, M. (2008). Spatially explicit maximum likelihood methods for capture-recapture studies. Biometrics, 64, 377–385. doi: 10.1111/j.1541-0420.2007.00927.x.

Chandler, R. B., & Clark, J. D. (2014). Spatially explicit integrated population models. Methods in Ecology and Evolution, 5, 1351–1360. doi: 10.1111/2041-210X.12153

Danancher, D., Labonne, J., Pradel, R., & Gaudin, P. (2004). Capture recapture estimates of space used in streams (CRE5U5) at the population scale: case study on Zingel asper (percid), a threatened species of the Rhône catchment. Canadian Journal of Fisheries and Aquatic Sciences, 61, 476–486. doi: 10.1139/f04-004

Ergon, T., & Gardner, B. (2014). Separating mortality and emigration: modelling space use, dispersal and survival with robust-design spatial capture-recapture data. Methods in Ecology and Evolution, 5, 1327–1336. doi: 10.1111/2041-210X.12133

Gardner, B., Reppucci, J., Lucherini, M., & Royle, J. A. (2010). Spatially explicit inference for open populations: estimating demographic parameters from camera-trap studies. Ecology, 91, 3376–3383. doi: 10.1890/09-0804.1

Gardner, B., Sollmann, R. Kuman, N. S., Jathanna, D., & Karanth, K. U. (2018). State-space and movement specification in open population spatial capture-recapture models. Ecology and Evolution, doi: 10.1002/ece3.4509

Helle P., Helle, T., & Linden, H. (1994). Capercaillie (*Tetrao urogallus*) lekking sites in fragmented Finnish forest landscape. Scandinavian Journal of Forest Research, 9, 386–396. doi: 10.1080/02827589409382856.

Jacob G., Debrunner R., Gugerli F., Schmid B., & Bollmann K. (2010) Field surveys of capercaillie (*Tetrao urogallus*) in the Swiss Alps underestimated local abundance of the species as revealed by genetic analyses of non-invasive samples. Conservation Genetics, 11, 33–44. doi: 10.1007/S10592-008-9794-8

Karanth, K. U., & Nichols, J. D. (1998). Estimation of tiger densities in India using photographic captures and recaptures. Ecology, 79, 2852–2862. doi: 10.1890/0012-9658(1998)079[2852:EOTDII]2.0.CO;2

Kéry, M., & Schaub, M. (2012). Bayesian population analysis using WinBUGS - A hierarchical perspective. Academic Press, Waltham, MA.

Mollet, P., Badilatti B., Bollmann, K., Graf, R. F., Hess, R., Jenny, H., Mulhauser, B., Perrenoud, A., Rudmann, R. Sachot, S., & Studer, J. (2003). Verbreitung und Bestand des Auerhuhns (*Tetrao urogallus*) in der Schweiz 2001 und ihre Veranderung im 19. und 20. Jahrhundert. Ornithologischer Beobachter, 100, 67–86.

Mollet, P., Kéry, M., Gardner, B., Pasinelli, G., & Royle, J. A. (2015). Estimating population size for capercaillie (*Tetrao urogallus* L.) with spatial capture-recapture models based on genotypes from one field sample. PLoS ONE 10(6): e0129020. doi:10.1371/journal.pone.0129020

O’Connell, A. F., Nichols, J. D., & Karanth, K.U. (2010). Camera traps in animal ecology - methods and analyses. Springer, New York.

Raabe, J. K., Gardner, B., & Hightower, J. E. (2013). A spatial capture-recapture model to estimate fish survival and location from linear continuous monitoring arrays. Canadian Journal of Fisheries and Aquatic Sciences, 71,120–130. doi: 10.1139/cjfas-2013-0198

Royle, J. A., & Dorazio, R. M. (2012). Parameter-expanded data augmentation for Bayesian analysis of capture-recapture models. Journal of Ornithology, 152, 521–537. doi: 10.1007/sl0336-010-0619-4

Royle, J. A., & Young, K. V. (2008). A hierarchical model for spatial capture-recapture data. Ecology, 89, 2281–2289. doi: 10.1890/07-0601.1

Royle, J. A., Chandler, R. B., Sollmann, R., & Gardner, B. (2014). Spatial capture-recapture. Academic Press. Waltham, MA, USA.

Royle, J. A., Fuller, A. K., & Sutherland, C. (2018). Unifying population and landscape ecology with spatial capture-recapture. Ecogaphy, 41(3), 444–456. doi: 10.1111/ecog.03170

Russell, R. E., Royle, J. A., Desimone, R., Schwartz, M. K., Edwards, V. L., Pilgrim, K. P., & Mckelvey, K. S. 2012. Estimating abundance of mountain lions from unstructured spatial sampling. Journal of Wildlife Management, 76, 1551–1561. doi: 10.1002/jwmg.412

Schaub, M., & Royle, J. A. (2014). Estimating true instead of apparent survival using spatial Cormack-Jolly-Seber models. Methods in Ecology and Evolution, 5, 1316–1326. doi: 10.1111/2041-210X.12134

Storch I. 2001. *Tetrao urogallus* Capercaillie. BWP Update 3:1–24.

Suter, W., Graf, R. F., & Hess, R. (2002). Capercaillie (*Tetrao urogallus*) and avian biodiversity: testing the umbrella-species concept. Conservation Biology, 16, 778–788. doi: 10.1046/j.1523-1739.2002.01129.x

Suwanrat, S., Ngoprasert, D., Sutherland, C., Suwanwaree, P. & Savini, T., 2015. Estimating density of secretive terrestrial birds (Siamese Fireback) in pristine and degraded forest using camera traps and distance sampling. Global Ecology and Conservation, 3, 596–606. doi: 10.1016/j.gecco.2015.01.010

Taberlet, P. & Luikart, G. 1999. Non-invasive genetic sampling and individual identification. Biological Journal of the Linnean Society, 68(1-2): 41–55. doi: doi.org/10.1111/j.1095-8312.1999.tb01157.x

## References

Bonin, A.; Bellemain, E.; Bronken Eidesen, P.; Pompanon, F.; Brochmann, C.; Taberlet, P. (2004): How to track and assess genotyping errors in population genetics studies. In: Molecular ecology 13 (11), S. 3261–3273. DOI: 10.1111/j.l365-294X.2004.02346.x.

Galpern, Paul; Manseau, Micheline; Hettinga, Peter; Smith, Karen; Wilson, Paul (2012): Allelematch. An R package for identifying unique multilocus genotypes where genotyping error and missing data may be present. In: Molecular ecology resources 12 (4), S. 771–778. DOI: 10.1111/j. 1755-0998.2012.03137.x.

Goudet J (2002) FSTAT, a program to estimate and test gene diversities and fixation indices, vers. 2.9.3.2. Available: http://www2.unil.ch/popgen/softwares/fstat.htm.

Guo SW, Thompson EA (1992) Performing the exact test of Hardy-Weinberg proportion for multiple alleles. Biometrics 48: 361–372. http://www.jstor.org/stable/2532296 PMID: 1637966

Kalinowski ST, Taper ML, Marshall TC (2007) Revising how the computer program CERVUS accommodates genotyping error increases success in paternity assignment. Molecular Ecology 16: 1099–1006. doi: 10.1111/j.l365-294X.2007.03089.x PMID: 17305863

Raymond M, Rousset F (1995) GenePop (version 1.2): population genetics software for exact tests and ecumenicism. Journal of Heredity 86: 248.

Rousset F (2008) Genepop’007: a complete reimplementation of the Genepop software for Windows and Linux. Molecular Ecology Resources 8:103–106. doi: 10.1111/j.l471-8286.2007.01931.x PMID: 21585727

Waits LP, Luikart G, Taberlet P (2001) Estimating the probability of identity among genotypes in natural populations: cautions and guidelines. Molecular Ecology 10: 249–256. doi: 10.1046/j.l365-294X.2001.01185.x PMID: 11251803

